# LabWeaver: reliable agentic data analysis grounded in laboratory knowledge

**DOI:** 10.64898/2026.10.28.762601

**Authors:** Li Li, Gabriela Fort, Jun Xu, Aik Choon Tan, Boyi Guo

## Abstract

Agentic systems powered by large language models are rapidly automating biological data analysis. However, their reliability remains questionable because they often overlook experimental context, such as disease domain and experimental protocol. Existing agentic systems organize context around individual users and projects, whereas research is organized around laboratories whose projects share experimental context. As a result, experimental context learned in one project remains unavailable to related projects. We introduce LabWeaver, an agentic data analysis framework with a laboratory-organized architecture of three tiers: laboratory, user and project. Further, LabWeaver distills experimental context into laboratory memory and skills, which are shared across users and projects after human-gated review. To isolate the contribution of experimental context, we analyzed a synovial sarcoma single-cell RNA-seq dataset in two sessions that differed only in whether laboratory memory and skills were available. The context-free session lost the fusion reporter at read alignment and misannotated the tumor cells. In contrast, the context-enabled session recovered reporter expression and resolved five fusion-anchored tumor cell states.

## Main

Agentic data analysis, empowered by rapid advancements in large language and reasoning models, has greatly lowered the technical barrier to perform complex bioinformatics tasks and data analysis. As a result, agentic data analysis and automation of scientific discovery are rapidly becoming an integral part of biomedical research.^1^ While agentic systems can automate data analysis with minimal to no oversight, the reliability of agentic data analysis remains questionable in many areas of computational biology, substantially lowering scientific confidence in generated results.^2,3^ There is thus an urgent need to improve the reliability of agentic data analysis and to ensure scientific rigor while accelerating the discovery process.

The design and execution of data analysis is a complex and iterative process, where subtle differences in analytical decisions can lead to conflicting conclusions.^4–6^ Experimental context, such as disease domain, model systems, and experimental design and protocol, provides a pivotal anchor for analytical decision making. Overlooking experimental context in data analysis often results in unreliable conclusions. Therefore, embedding experimental context into an agentic system is critical to improve the reliability of agentic data analysis.

## Inherent misalignment between existing agentic systems and research practice

Existing agentic systems are built around individual projects. This user-project architecture limits learning by preventing the system from transferring and leveraging experimental context across projects. (**Figure 1a**) Context enters the system only when users supply it via prompts, and it stays inside that project. As a result, context learned in one project is unavailable to other contextually similar projects. Additional strategies have been developed to strengthen these systems, but they still largely rely on general-purpose pipelines and underutilize the experimental context specific to the project, only modestly improving the reliability of data analysis.^2,3^ For example, retrieval-augmented generation frameworks leverage curated databases, knowledge graphs, and published literature to augment analytical decision making.^7,8^ However, the analytical decisions that shape a result are often incompletely specified in published methods and are not specifically tailored to a given research program or domain.^9^ Self-evolving agents accumulate tools and skills across the tasks they complete;^10,11^ however, they learn only from their own past runs rather than across related projects or domains and thus lack appropriate experimental context.

**Figure 1.**
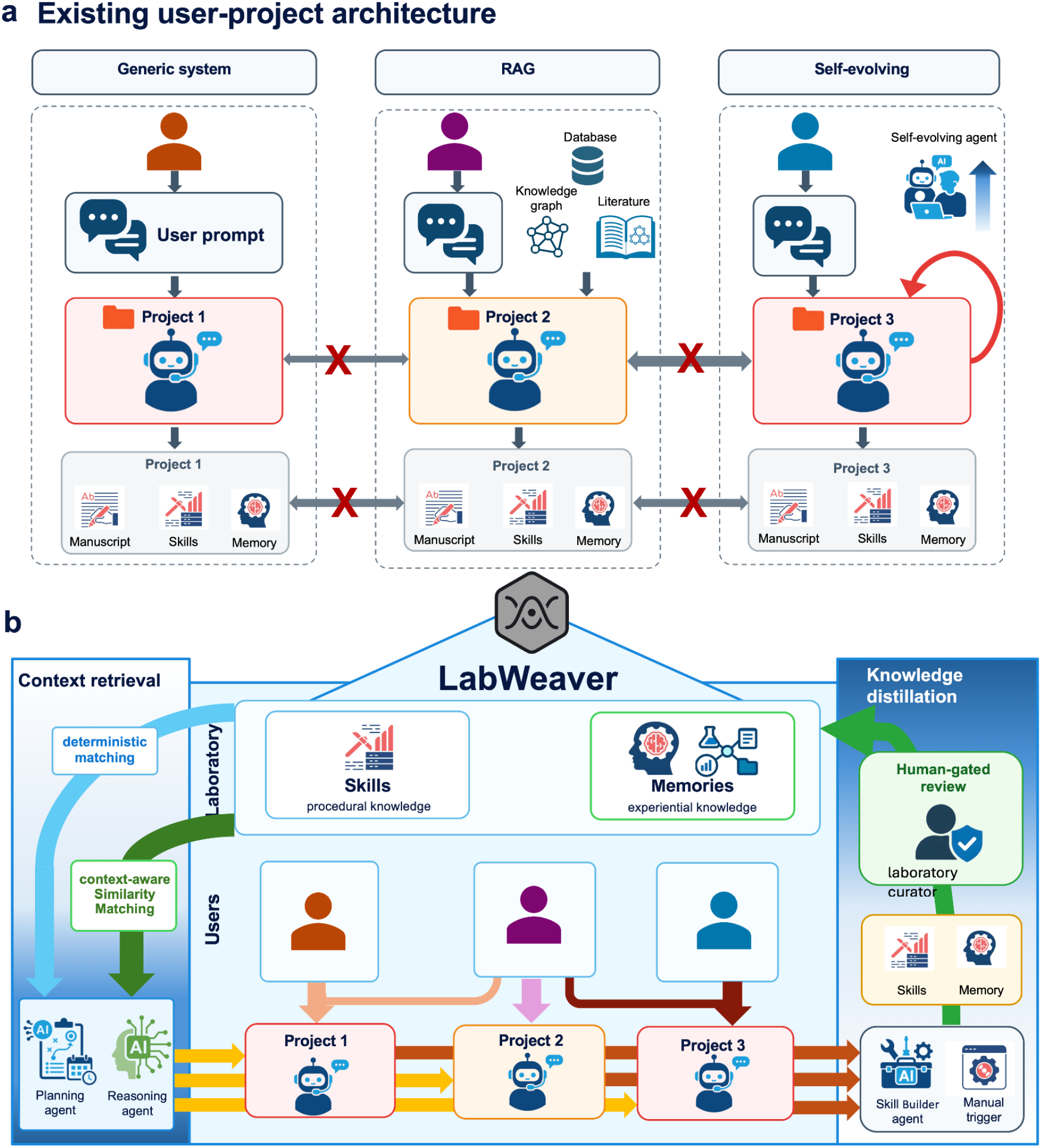
Misalignment of existing user-project agentic architectures with laboratory-based research practice can be addressed by the proposed LabWeaver framework. (**a**) Existing agentic systems are organized around isolated user-project units. In a general-purpose agent, experimental context is supplied primarily through the user prompt and remains confined to the project, optionally augmented by retrieval-augmented generation (RAG) or self-evolution. (**b**) LabWeaver introduces a laboratory-organized architecture with three tiers (laboratory, user and project) and knowledge distillation and retrieval mechanisms built to store and populate context along the three-tier architecture.

In reality, research is organized around laboratories. A research laboratory is built around a shared research question, and its projects are typically contextually related. For example, successive studies investigate related disease mechanisms; data are generated from the same model organisms following the same experimental protocols with similar platforms and sequencing depths. This contextual information informs analytical decision making, which needs to be fine-tuned for different laboratories who follow different methods and experimental protocols. Moreover, domain-specific analytical expertise evolves across projects through a record of trial and error that is seldom written down and never published.^12,13^ The misalignment between laboratory-organized research practice and the user-project architecture of current agentic systems creates a substantial gap that prevents contextual information from reaching the analysis, and hence impedes reliable agentic data analysis.

## LabWeaver, a laboratory-organized architecture

Here we introduce LabWeaver, an agentic data analysis framework organized around the laboratory rather than the isolated project. (**Figure 1b**) The key innovations of LabWeaver are twofold: 1) a laboratory-organized architecture with three tiers, i.e., laboratory, user and project; and 2) knowledge distillation and retrieval mechanisms built to store and populate context along the three-tier architecture. These innovations also support a continuous learning cycle in which laboratory experience informs the evolution of laboratory practice.

LabWeaver extends the existing user-project architecture by adding a laboratory tier above it, within which users and their projects are hosted. This laboratory-user-project architecture makes experimental context a property of the laboratory, and enables context to be learned and retrieved across projects and users. Moreover, by introducing the laboratory tier, the institutional knowledge accumulated through individual users and projects is preserved when a user leaves the laboratory. At the user tier, LabWeaver grants a designated subset of users, e.g., the head of the laboratory, as *laboratory curators*. Laboratory curators govern knowledge distillation into the laboratory tier and retain the authority to revise or retract entries later found to be incorrect. LabWeaver requires all context to pass a human-gated review by laboratory curators before it is released for laboratory-wide use. The remaining tiers follow the user-project architecture, preserving user- and project-specific context.

To account for experimental context in analytical processes, LabWeaver digitizes context as *skills* and *memories* to store *procedural* and *experiential knowledge*, respectively. Skills, in the form of markdown files, preserve validated and reusable computational protocols; memories, in the form of knowledge graphs and relational databases, organize experimental records, unpublished data, and tacit laboratory expertise, such as domain-guided analytical and scientific reasoning. While retaining user- and project-tier skills and memories, LabWeaver introduces a laboratory-tier context library that enables skills and memories to be shared across projects. In a project, laboratory-tier skills and memories are accessed by agents through different mechanisms. Skills are retrieved during the planning stage, though they can also be manually invoked. The planning agent selects skills by deterministic matching to tasks, based on analytical intention and keywords such as analysis type, data modality, and workflow requirements. In contrast, memories are accessible throughout the project by both planning and reasoning agents. These agents retrieve memories using a *context-aware similarity matching* strategy by comparing a current task with both the content and contextual metadata of stored memories and favoring memories that are semantically relevant and arise from a similar research context. These innovations in LabWeaver enable efficient sharing and integration of experimental context in analysis, hence improving the reliability of data analysis.

To facilitate context accumulation within the laboratory, LabWeaver introduces a novel knowledge distillation mechanism through which a laboratory’s domain expertise develops as its work proceeds. Skills are created and refined by a Skill Builder agent and stored in a structured skill format. A memory is captured when a researcher determines that the current session produced a successful or reusable outcome.

Critically, neither skills nor memories enter the shared laboratory-tier context library automatically. LabWeaver requires all context to pass a human-gated review process by laboratory curators before it is released for laboratory-wide use. This separation of laboratory-wide context from individual- and project-specific ones provides a controlled path to become trusted institutional knowledge. In addition, the learning of experimental context can also take the form of reproducing published analyses and constructing pipelines de novo from public data (see **Case study 2**). Overall, this mechanism enables LabWeaver to preserve experiential knowledge and capture comprehensive workflow.

### Case study 1: experimental context recovers fusion reporter expression and resolves synovial sarcoma cell states

To illustrate how experimental context can improve agentic data analysis, we applied LabWeaver to an in-house single-cell RNA sequencing (scRNA-seq) dataset to investigate the tumorigenesis of synovial sarcoma, an aggressive soft-tissue cancer.^14^(**Figure 2a**) Briefly, scRNA-seq data were generated from a mouse model^15^ carrying the *SS18::SSX2* fusion that drives synovial sarcoma, the product of a recurrent t(X;18) translocation. *IRES-EGFP*, a reporter transcribed together with the fusion, was designed to identify which cell population harbored the *SS18::SSX2* fusion. To isolate the contribution of experimental context, we instructed LabWeaver to autonomously structure and execute data analysis pipelines in two isolated computation sessions that differed only in whether laboratory memory and skills were available. We refer to these sessions as *context-enabled* and *context-free*, respectively. Of note, LabWeaver without experimental context reverts to the underlying agentic system, and hence, we used it to approximate how the user-project architecture would perform. Although both sessions used the same agent and identical inputs, they produced substantially different analysis outcomes. In the context-free session, EGFP was not leveraged as a tumor-specific marker and the tumor compartment collapsed into a single mislabeled population. (**Figure 2b**) In contrast, in the context-enabled session, the same cells were resolved into five fusion-anchored states. (**Figure 2c,d**)

**Figure 2.**
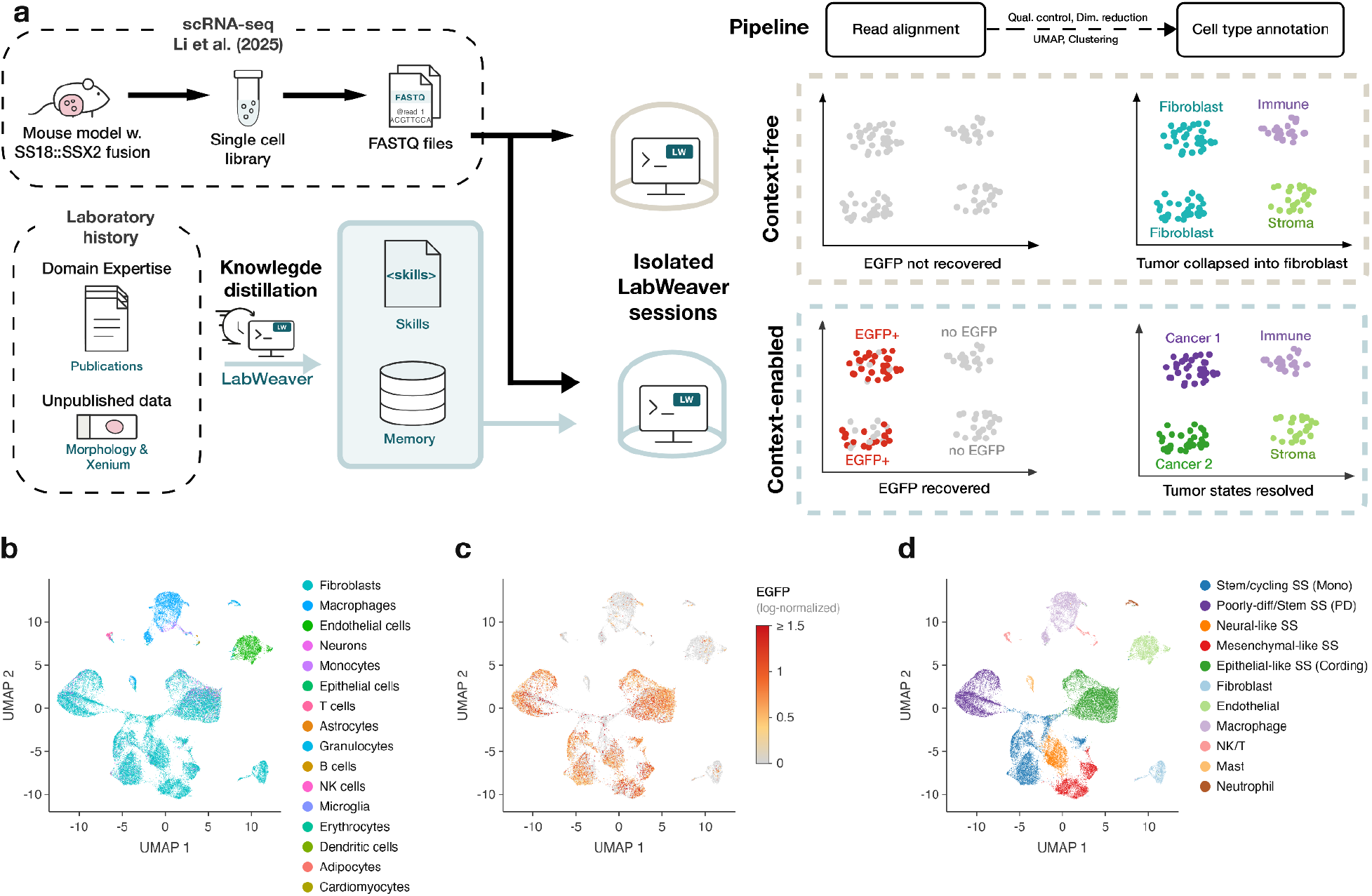
Laboratory memory and skills resolve SS18::SSX2 reporter expression and synovial sarcoma cell states. (**a**) The schematic describing the *in silico* experiment that examines LabWeaver’s ability to leverage experimental context to improve single-cell RNA-seq data analysis. (**b-d**) Annotated cells from context-free and context-enabled sessions are mapped in the same UMAP. (**b**) Cell type annotations from the context-free session misaligns tumor cells to fibroblasts, neurons and epithelial cells. (**c**) Log-normalized EGFP expression produced from the context-enabled session. (**d**) Cell type annotations from the context-enabled session resolves tumor cells to cell states leveraging marker panels curated from prior Xenium and histological analyses in the laboratory library.

To understand what produced the discrepancy, we dissected the pipeline each session built. Surprisingly, the two sessions diverged from the very first step of the pipeline, read alignment. While both sessions used the same tool, Cell Ranger (v9.0.1), the context-free session aligned all reads to a standard mouse genome reference (GRCm39). Because that reference contains no EGFP contig, reads spanning the reporter failed to align and the transcript was absent from the count matrix, although it was present in the sequencing library. In contrast, the context-enabled session recognized the experimental design, retrieved the laboratory’s reporter-augmented reference from memory, and aligned against it without being asked, recovering per-cell EGFP counts. (**Figure 2c**) As EGFP plays an irreplaceable role in this biological investigation, this divergence propagates through the analysis pipeline and subsequent biological interpretation of the findings.

The two sessions also provided drastically different biological interpretations of the surveyed cell population. The context-free session applied a general-purpose cell-type reference (SingleR v1.0) that lacks synovial sarcoma representation. As a result, the majority of tumor cells were mistakenly annotated as transcriptomically similar but biologically inaccurate cell archetypes, specifically 29,813 as fibroblasts 2,458 as neurons, 233 as epithelial cells and 171 as astrocytes. (**Figure 2b**) This misannotation likely reflects the loss of EGFP, the tumor-cell marker, at the read alignment step earlier. In contrast, the context-enabled session drew on marker panels stored in the laboratory memory. As a result, five synovial sarcoma states, namely epithelial-like/cording, stem/cycling, poorly differentiated/stem, mesenchymal-like, and neural-like, were resolved in the context-enabled session. (**Figure 2d**)

## Case study 2: LabWeaver reproduces scientific investigation through publication

To demonstrate the ability of LabWeaver to reproduce published scRNAseq results, we used the raw data and analysis methods from a recent study of lung adenocarcinoma^16^. This study used a genetically engineered mouse model to examine how deletion of the transcription factor *Hnf4a* alters tumor cell identity. The experimental design included control and *Hnf4a*-deleted tumors, with paired samples either left intact or depleted of stromal and immune cells to enrich for tumor cells.

To establish the analysis environment, we uploaded Cell Ranger output files obtained from GEO (GSE281964) as referenced in the manuscript. LabWeaver was provided with the sample identities and published methods and was asked to reproduce the published analysis.

LabWeaver successfully implemented the major steps described, comprising a standard Seurat-based workflow including quality control (QC), normalization, dimensionality reduction, clustering, UMAP visualization, cell type annotation, and differential gene expression analysis. Importantly, the tool adhered to key implementation details necessary for reproducibility, such as applying the same QC thresholds, clustering resolution, and dimensionality used for downstream analyses. The reproduced analysis closely matched the published results. The initial UMAP embedding separated samples according to stromal and immune cell depletion, as reported in the original study (**Figure 3A-B**). Using established marker genes, LabWeaver distinguished tumor cells from stromal and immune populations and reclustered focusing only on the tumor cell compartment. The resulting tumor cell structure closely resembled the published embeddings and clusters (**Figure 3C-D**). Most importantly, LabWeaver reproduced one of the central biological results from the study. Differential expression between genotype-enriched tumor cell clusters showed that *Hnf4a* deletion was associated with a loss of gastric- and liver-like transcriptional programs and increased expression of lung alveolar programs (**Figure 3E**). Thus, LabWeaver reproduced both the computational structure of the analysis given the methods as context and the principal biological interpretation from this published experiment. Together, this case study demonstrates that LabWeaver can translate published methods and raw data into a reproducible scRNA-seq workflow, and hence is capable of learning context through study replication.

**Figure 3.**
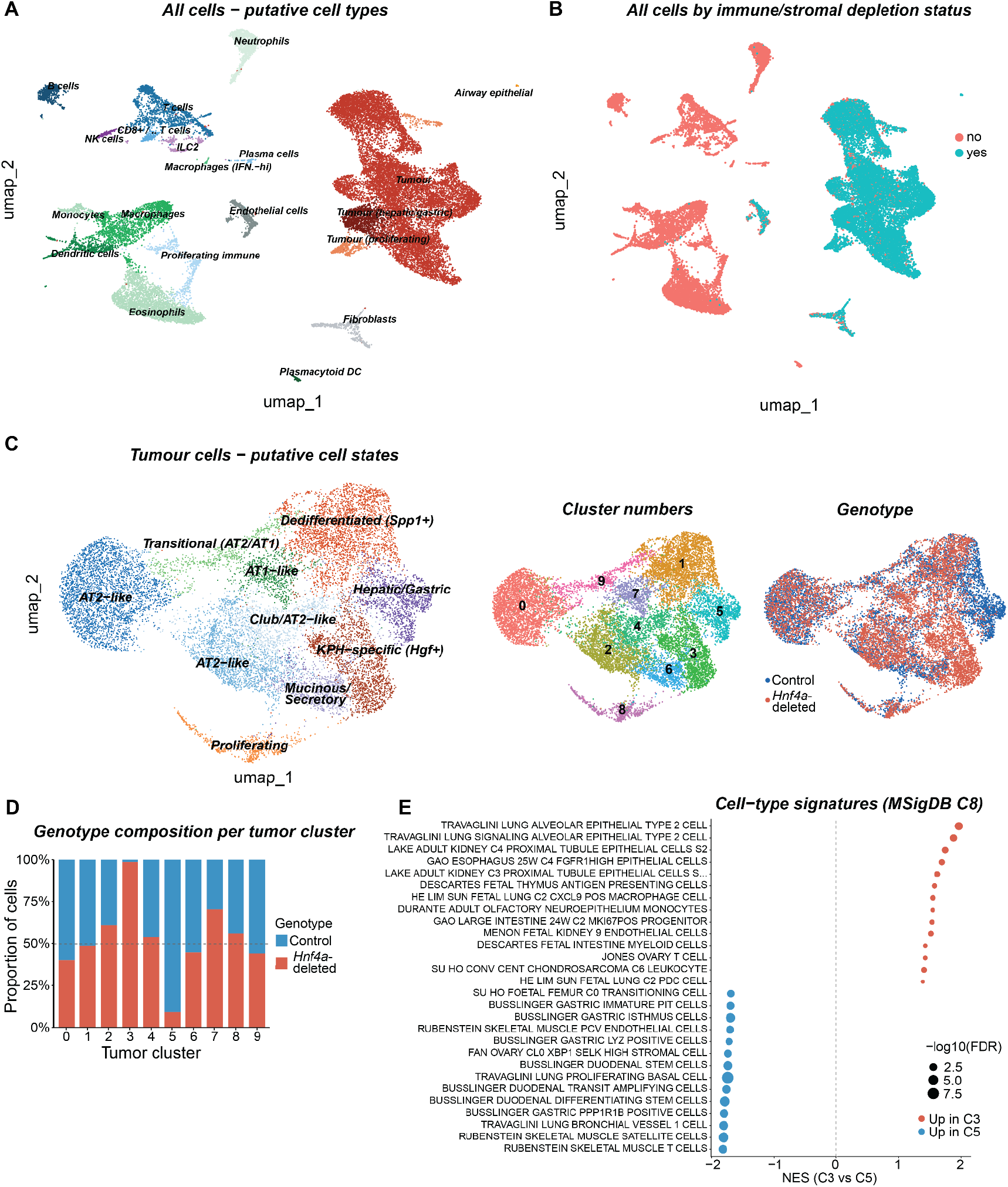
LabWeaver reproduces published scRNA-seq analysis. (A) UMAP of all cells colored by inferred cell type based on marker gene expression. (B) UMAP colored by stromal/immune depletion status. (C) UMAP of tumor cells after subsetting and reclustering. Clusters are annotated by putative tumor cell states based on marker gene expression, cluster number, or genotype. (D) Genotype composition across tumor cell clusters. (E) Gene set enrichment analysis (MSigDB C8) comparing control-enriched versus *Hnf4a*-deletion-enriched clusters.

## Discussion

We introduced LabWeaver, an agentic data analysis framework organized around the laboratory rather than the isolated project. LabWeaver addresses a gap that existing agentic frameworks leave open, how to incorporate experimental context into agentic analysis, by introducing a three-tier architecture and innovative knowledge distillation and retrieval mechanisms. Applied to a synovial sarcoma scRNA-seq study, LabWeaver delivers more biologically informative analysis and interpretation of the data by leveraging the experimental context of the study.

The case study above demonstrates that experimental context improves the reliability of agentic data analysis, which is the central contribution of LabWeaver to biological data analysis. It is a natural future step to evaluate the impact of experimental context on agentic data analysis across molecular assays and organ systems, where we anticipate heterogeneous performances. However, such systematic evaluations of context-enabled agentic platforms will require designs that are different from existing benchmarking frameworks that focus on evaluating the underlying language models rather than the contribution of experimental context. This need for additional benchmark frameworks reflects a gap between agentic system development and data analysis practice, which often results in overoptimism and overconfidence in real-world applications that could fuel the replication crisis in science if left unaddressed. We therefore consider the design of benchmark studies for context-dependent analysis an important direction for the development of agentic data analysis.

Beyond data analysis, LabWeaver’s laboratory-organized architecture provides a natural platform to support practical developments for laboratories and research ecosystems. Knowledge preservation in academic laboratories is challenged by the transient nature of trainee appointments. LabWeaver is capable of digitizing and recording tacit knowledge and members’ experience in the laboratory-tier context library. Reciprocally, the context library facilitates the onboarding of new members and intra- and inter-laboratory collaborations by lowering communication and logistical burdens. Furthermore, LabWeaver provides the natural infrastructure to future-proof data for their AI readiness.^17^ Standard-binding data management and stewardship and comprehensive workflow capture can be easily automated in LabWeaver. Future integration with specialized agents^18,19^ will facilitate education for trainees. Overall, LabWeaver presents a unique opportunity to address the missing data challenge faced by intelligent scientific instruments^20^, bringing expert-level decision-making and troubleshooting skills to autonomous systems.

## Back Matter

### Code and Data availability

LabWeaver is publicly available at (https://github.com/lili-8477/labweaver). To reproduce the case study, containerized LabWeaver sessions are available at https://labweaver.ai/app?auto=true&nats=wss%3A%2F%2Flabweaver.ai%2Fws%2F using service id for context-enabled (1240201e2179d9fd2473f51a88b5d862916f7a203251e96555a6e1a0eabef1aa) and context-free (68bcef55d23993db582fb90dc826fa83263d8794d6221e1ccd24a826d96340a8).

## Acknowledgements, Funding, Authorship Contributions

## Acknowledgements

The authors acknowledge the use of AI-assisted coding tools to support the implementation of LabWeaver. We thank Dr. Eric Snyder for providing feedback on the biological validity of results produced by LabWeaver.

## Author Contributions

Conceptualization: LL, ACT, BG Methodology: LL

Software: LL, JX Formal analysis: LL, GF

Investigation: LL, GF, BG Data Curation: LL

Writing - Original Draft: LL, GF

Writing - Review & Editing: LL, GF, JX, ACT, BG Visualization: LL, GF, BG

Supervision: ACT, BG Project administration: BG

Funding acquisition: ACT, BG

